# Spatially-constrained growth enhances conversional meltdown

**DOI:** 10.1101/027292

**Authors:** Maxim O. Lavrentovich, Mary E. Wahl, David R. Nelson, Andrew W. Murray

## Abstract

Cells that mutate or commit to a specialized function (differentiate) often undergo conversions that are effectively irreversible. Slowed growth of converted cells can act as a form of selection, balancing unidirectional conversion to maintain both cell types at a steady-state ratio. However, when one-way conversion is insufficiently counterbalanced by selection, the original cell type will ultimately be lost, often with negative impacts on the population’s overall fitness. The critical balance between selection and conversion needed for preservation of unconverted cells and the steady-state ratio between cell types depends on the spatial circumstances under which cells proliferate. We present experimental data on a yeast strain engineered to undergo irreversible conversion: this synthetic system permits cell type-specific fluorescent labeling and exogenous variation of the relative growth and conversion rates. We fnd that populations confined to grow on a flat agar surface are more susceptible than their well-mixed counterparts to fitness loss via a conversion-induced “meltdown.” We then present analytical predictions for growth in several biologically-relevant geometries – well-mixed liquid media, radially-expanding two-dimensional colonies, and linear fronts in two dimensions – by employing analogies to the directed percolation transition from non-equilibrium statistical physics. These simplified theories are consistent with the experimental results.

## Introduction

Irreversible change is an important aspect of both development (1) and evolution (2). Many mature tissues retain stem cells that replenish specialized cells lost to damage or aging. Proliferation balanced by irreversible differentiation can maintain stem and specialized cells in a dynamic steady-state (3), but an imbalance between these forces can eliminate the stem cell population, with dire health consequences (4). Like differentiation, harmful mutations can be effectively irreversible; natural selection can check their spread if the mutants reproduce more slowly, but if the mutation rate is too great or selection too weak, these mutations can fix permanently. Such a mutational meltdown is known as Muller’s ratchet in the population genetics literature (5, 6). We will employ the generic term “conversional meltdown” to describe the loss of an unconverted cell type due to an unfavorable balance between mutation and selection, differentiation and proliferation, and, more generally, any form of irreversible conversion and differential growth. The abrupt shift from maintenance to extinction of the unconverted cell type as conversion rate increases is analogous to the well-studied directed percolation phase transition in statistical physics (7–9).

Though most analyses of this important phase transition have focussed on well-mixed populations, spatial structure can play a crucial role (8, 10, 11). Here, we investigate conversional meltdown for one-dimensional growth without subsequent migration, a geometry relevant in natural circumstances such as population expansions and growth of the plant meristem, as well as in experimentally-tractable systems such as microbial range expansions (12, 13). Yeast (13) and immotile bacteria (12) on Petri dishes grow in colonies that remain relatively flat, proliferating primarily at the edges (14). Due to the small effective populations which compete to divide into virgin territory, the thin region of dividing cells at the frontier can be treated as a one-dimensional population. Nutrient depletion in the colony core preserves the colony interior, which reflects the past history of such populations: the balance between cell types can be studied using fluorescence detection techniques. When a particular cell type has locally fixed at the colony frontier, its descendants form a “sector” as shown in blue in Fig. 1(a). The geometric properties of the spatial sectors reflect the underlying evolutionary dynamics: for example, the sector opening angle *θ* provides an estimate of the selective advantage of cells in the sector relative to their neighbors (13, 14).

**Figure 1.**
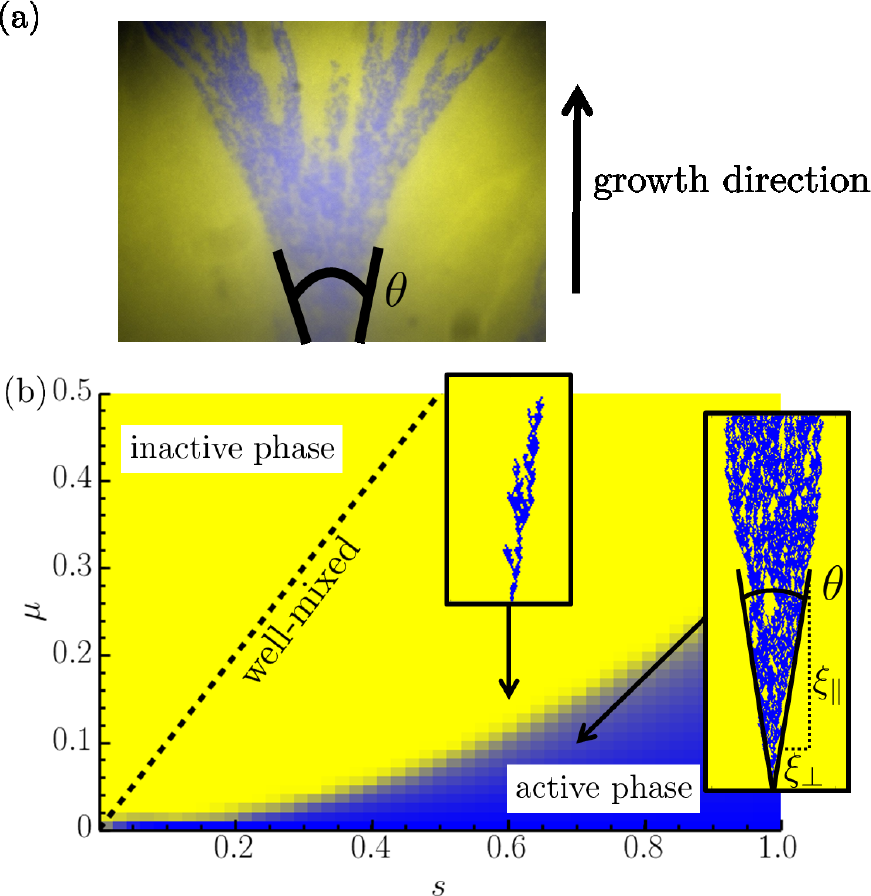
(a) A micrograph of the edge of a two-dimensional budding yeast *S. cerevisiae* cell colony grown on a Petri dish. In this linear range expansion (characterized by an conversion rate *μ* and a selective advantage ***s*** of blue cells over yellow cells), the blue unconverted cells form a spatial sector with an opening angle ***θ*** marked with overlaid black lines. With each cell division, the blue cells enjoying a selective advantage ***s*** convert to the yellow ones at a rate μ,which creates yellow patches within the blue sector. The growth, or time-like, direction is indicated. (b) A phase diagram indicating where theory predicts the eventual extinction of the blue strain as a function of its selective advantage ***s*** and conversion rate *μ* for a linear range expansion. In the yellow “inactive” phase, a genetic sector formed by a blue cell always dies out, leading to a fully converted population. In the blue “active” phase, there is a non-zero probability of forming a surviving cluster, leading to a population with conversion occurring stochastically. The transition line for a well-mixed population is also shown for comparison (dashed line). In the well-mixed case, the active phase forms a much larger region in the (***μ, s***) -plane. The insets show examples of simulated sectors. Note the resemblance between the sector in the active phase and the experimental sector in part (a).

This investigation focuses on the effect of spatial population structure on the conversional meltdown phase transition. We perform *in vivo* experiments, complemented with analytical and simulation-based approaches, which show the striking effects of spatial structure on evolutionary dynamics. We employ a strain of budding yeast engineered to undergo irreversible conversions with independently tunable frequency and fitness cost to study population dynamics in well-mixed liquid media, as well as microbial range expansions on Petri dishes. We find that the spatial distribution of the cells qualitatively changes the dynamics. Only adjacent individuals in spatially distributed populations compete, and the local effective population size is thus small relative to the total population. The small number of competing individuals amplifies the important of number fluctuations, i.e., genetic drift. We will show through experiments, simulation, and theory that this enhanced genetic drift significantly favors extinction, relative to well-mixed situations. This enhancement of extinction may have important consequences for diverse processes including tissue renewal (3), meristematic growth (15), and mutation-selection balance (16), since the relative proliferation of the unconverted strain must occur faster in a spatially distributed population than expected from experiments on well-mixed populations to prevent extinction of the unconverted population.

Crucially, the extinction transition we study here is distinct from extinction due to neutral competition dynamics. For example, previous studies of stem cells in intestinal crypts (3, 10, 17) found that different stem cell clones may compete until a single clone takes over the whole population, while the other clones go extinct. These previous studies found that the clones are neutral with respect to each other, so that any one of them may take over. However, apart from this competition, the clones also terminally differentiate into other cell types. The evolutionary dynamics of this differentiation process is not expected to be neutral, as the differentiated cells may reproduce more slowly and suffer a selective disadvantage. Moreover, even when the selection is weak, the associated extinction transition is of a different type from neutral competition: its dynamical scaling laws are governed by spatial mutation-selection balance and not by genetic drift alone. Since different cell types generically have different growth rates, we expect that our theory and experimental results describe features of extinction transitions in a broad range of biological systems.

## Materials and Methods

Microbes such as the budding yeast, *Saccharomyces cerevisiae*, are easily cultured in both test tubes and on Petri dishes. This makes them excellent candidates for comparing well-mixed and two-dimensional spatial dynamics. Construction of a yeast strain which undergoes irreversible conversion events with exogenously-tunable conversion rates and fitness cost was described in Ref. (18). Briefly, a *S. cerevisiae* strain was genetically engineered to lose a cycloheximide resistant ribosomal protein coding sequence via excision of a fragment of DNA by site-specific recombination. The activity of Cre, the site-specific recombinase, was controlled by varying the concentration of *β*-estradiol in the medium as described by Lindstrom et al. (19). This irreversible conversion event occurs once per cell division (during mitotic exit) with a probability *μ*, which we will call the conversion or mutation rate (per division). The probability **μ** depends on the ambient *β*-estradiol concentration. The cycloheximide resistant sequence (the *cyh2^r^* allele of the ribosomal protein L28 (20)) confers a measurable selective advantage for the unconverted strain relative to the converted strain when the strains are grown in the presence of cycloheximide. The precise selection coefficient *s* ≥ 0 associated with this advantage is tunable by varying the cycloheximide concentration in the medium. Both the conversion rate **μ** and selection coefficient *s* can be directly measured in well-mixed media and tuned over more than an order of magnitude by selecting appropriate *β*-estradiol and cycloheximide concentrations. Since these compounds are not consumed by the cells and diffuse readily through agar, and because yeast colonies are not particularly thick, these measurements also determine **μ** and *s* for populations grown on agar media.

To measure the fraction of converted versus unconverted cells in the population over time, we labeled the two cell types with fluorescent markers. Specifically, the coding sequence for the fluorescent protein mCherry is excised along with *cyh2^r^* via the Cre-mediated recombination. After the recombination event, an mCitrine fluorescent protein is expressed, instead. This set-up allows us to monitor the unconverted and converted cells using two different fluorescence channels. Throughout this manuscript, we have chosen to color the unconverted, mCherry-expressing cells blue and the converted, mCitrine-expressing cells yellow (see Fig. 1(a), for an example).

To visualize the conversional meltdown (i.e., the directed percolation transition), we produced linear range expansions on 1% agar media with judiciously-chosen *β*-estradiol and cycloheximide concentrations. To initiate the expansion, a thin strip of Whatman filter paper was submerged in a well-mixed liquid containing unconverted and converted cells, then placed in the center of the Petri dish; the linear colonies were then imaged after seven days’ growth (corresponding to about a 1 cm advancement of the colony front) at 30 °C. The ratio of unconverted to converted cells in the inoculum was chosen to be small enough so that resulting sectors of unconverted cells would typically be sufficiently separated for easy analysis. Fig. 2 displays representative images for colonies grown in a variety of agar media, the concentrations of *β*-estradiol ([*β*-est]) and cycloheximide ([CHX]) used in each, and the corresponding *μ* and *s* values (as determined in well-mixed media at the same [CHX] and [*β*-est]). The different preparations influence the range expansion dynamics: we see that either increasing *β*-estradiol concentrations or decreasing cycloheximide will yield smaller blue sectors in Fig. 2, indicating an approach to extinction of the unconverted blue strain. We will also consider range expansions in which we place a droplet of the yeast cell solution at the center of the Petri dish, which then forms a circular colony that spreads out radially.

**Figure 2.**
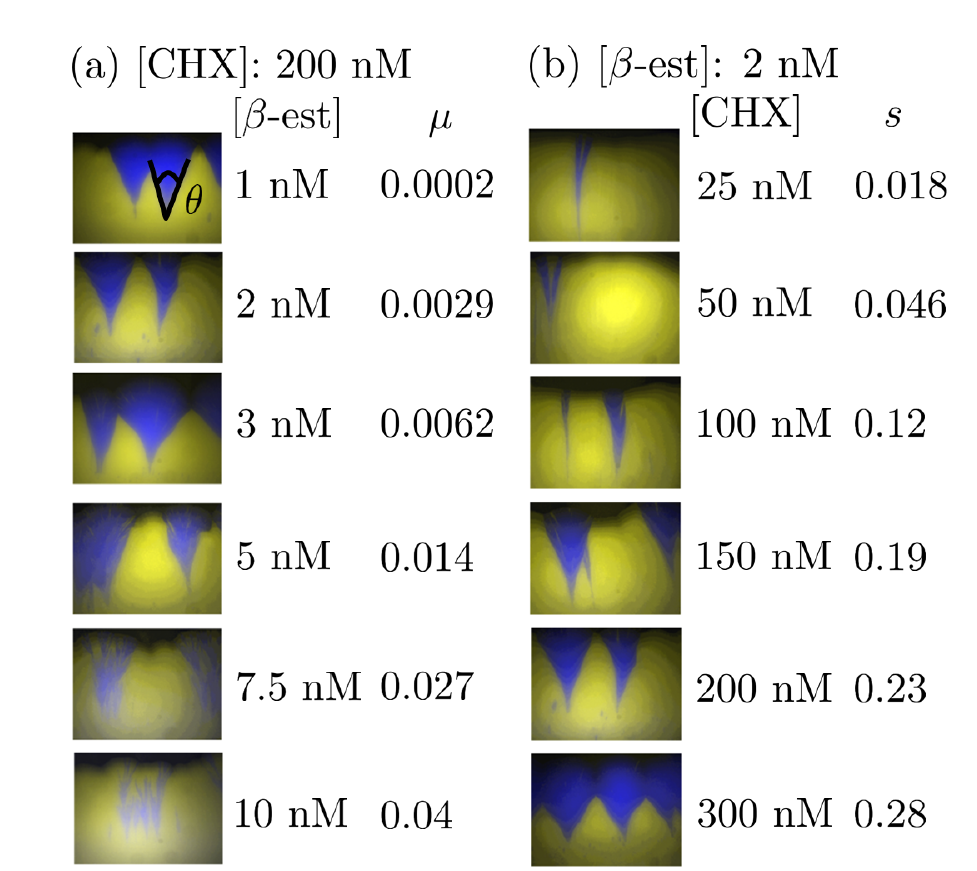
Frontiers of linear range expansions under different growth conditions. In (a), the *β*-estradiol concentration in the agar is varied with a fixed cycloheximide concentration. The corresponding conversion rates *μ* are indicated. In the top-most panel, we indicate an opening sector angle. In (b), the cycloheximide concentration is varied instead, tuning the selective advantage *s* of the blue strain over the yellow over a broad range. Note that the sector angles get smaller as we either increase *μ* or decrease *s* to approach the directed percolation (conversional meltdown) transition.

Thus, we are able to manipulate *μ* and *s* in experiment by varying the concentrations of *β*-estradiol and cycloheximide, respectively, in either the nutrient medium for well-mixed populations grown in test-tube, or in the agar for populations grown on plates. Note that it is possible to vary *μ* and *s* over a large range, covering values in the simulated phase diagram in Fig. 1(b): in particular, we are able to tune through the line separating the active and inactive phases and see extinction of the unconverted strain. We will now present experimental results on populations near this transition line.

## Results and Discussion

### Experimental Results

We first compare the steady-state concentration of unconverted blue cells in well-mixed populations and two-dimensional range expansions as a function of the mutation rate *μ* and the blue cell selective advantage *s*. We expect that if *μ* is large enough compared to *s*, the fit blue strain will be unable to survive in the population at long times, and the average fraction of blue cells 〈*f*〉 will eventually decay to zero. However, if *μ* is small, the fraction will approach some non-zero steady-state value *f_∞_*. We estimate this value in the well-mixed populations by measuring the fraction of mCherry-expressing unconverted cells by flow cytometry after enough generations to achieve a steady-state (approximately 40), or until the unconverted fraction is no longer measurable (18). Similarly, we estimate the fraction of unconverted cells in colonies at steady-state by collecting cells from the very edge of circular colonies after five days’ growth with a pipette tip and performing flow cytometry. The population frontier inflates in the circular range expansions, which has consequences for the dynamics. However, the steady-state fraction *f_∞_* is insensitive to this change in geometry (9).

The experimental results in Fig. 3 illustrate the striking effect of spatial fluctuations on the transition to extinction: compared to the well-mixed case, there is a significantly smaller section of the (*μ, s*) space that yields a non-zero steady-state fraction of unconverted cells in the population. The theoretical predictions for the phase boundaries (described in detail in the next section) are consistent with the experiment; We find *μ* ≈ *s* for the well-mixed population and *μ* ≈*A s^2^* for the populations grown on Petri dishes, where *A* ≈ 1.5 is a fitting parameter.

**Figure 3.**
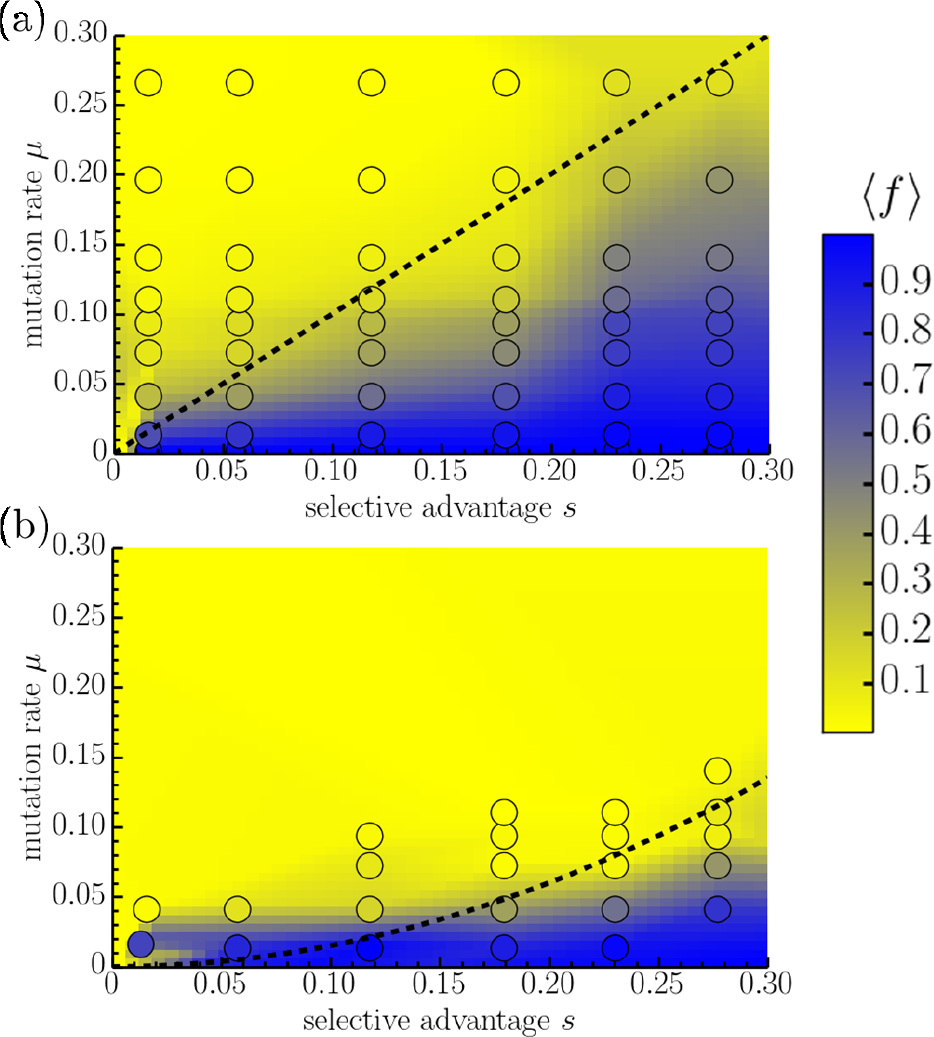
The average steady-state fraction of unconverted cells 〈f〉 at long times in (a) well-mixed populations cultured in a test-tube and in (b) two-dimensional range expansions. The concentration for the range expansions was measured by sampling cells at the edge of a circular colony after fve days of growth. The circles are the collected data points, which are used to make the interpolated color density plot in the background. The dashed lines are the theoretical predictions of the phase transition lines (see section titled Theory and Simulation). In (a), we expect that the transition occurs around *μ ≈ s*. In (b), we fnd a significantly different line shape, consistent with *μ* œ *As^2^*, with *A* ≈ 1.5 as the single parameter ft to the data.

It is also interesting to study the opening angle *θ* formed by the sectors in linear range expansions as we approach the phase transition line. The measured opening angles as a function of Δ are shown in Fig. 4. The values are collected by approximating the opening sector angle from images of the colony edges and averaging over many sectors. The error bars are calculated from the standard deviations of the sector angle measurements used to compute the averages. Growth conditions corresponding to many different values of *μ* and *s* were used, as illustrated in the inset of Fig. 4. The experiments are consistent with the theoretical prediction described in detail in the next section, except for small Δ. Note that in this regime, the sector angles are quite small and it is difficult to resolve them in the range expansion images. It would be interesting to study this regime in more detail in the future with better-resolved sector angle images to see if the directed percolation theory describes the experiments, or if a more sophisticated theory is necessary.

**Figure 4.**
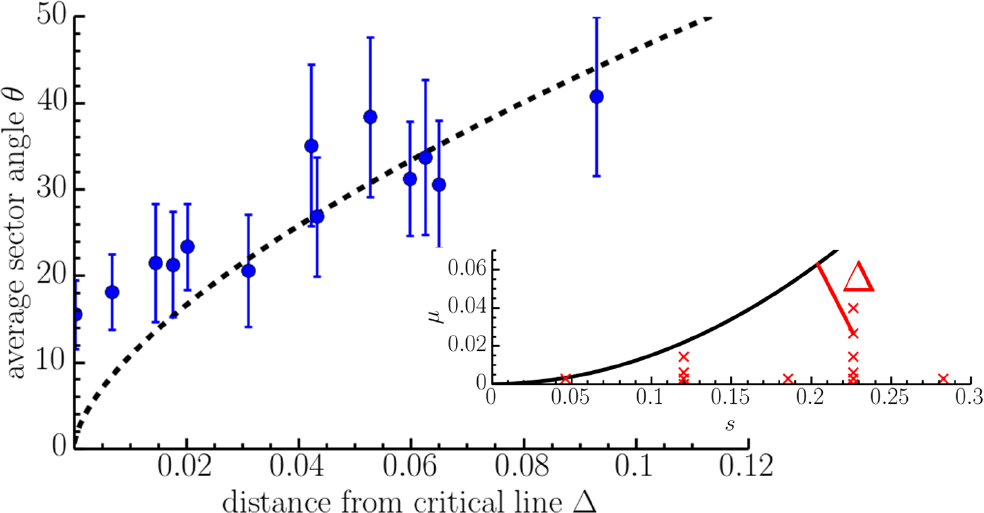
Measured average opening angle of sectors formed in experimental linear range expansions as a function of the shortest distance Δ from the critical line found in Fig. 3(b). The dashed line shows the ft to the expected directed percolation power law behavior discussed in the section titled Theory and Simulation (see Eq. 4). Inset: The black lines show the position of the transition, as determined in Fig. 3(b). The red crosses show the (*μ*, *s*) coordinates of all the growth conditions used to grow the colonies in the experiment. The distance Δ is also shown for one of these points with a solid red line.

### Theory and Simulation

We will now develop a theory for the observed experimental results based on the well-studied directed percolation phase transition (7). We begin with some approximations: As the yeast cell colony spreads across the agar plate, the evolutionary dynamics of interest occur at the frontier where cells settle virgin territory. Because yeast cells have low motility, cells which are even a few cell diameters behind the advancing population front may not be able to contribute to the population at the frontier, even if they continue to divide. Hence, we expect that the effective population of cells at the frontier which competes to divide into new territory is small. Thus, we focus our theoretical analysis on the population of cells living on a thin region at the colony frontier. This assumption is consistent with a previous study of two-dimensional colonies of mutualistic yeast, which also have relatively small effective population densities (21). Then, provided the yeast colony experiences a strong effective surface tension that forces the colony boundary to remain approximately circular, we may consider the dynamics along a uniform, effectively one-dimensional flat front. This geometry is consistent with microscopic observations of the yeast colony frontier (13). Note that rough fronts can significantly modify the nature of the extinction transition (22).

Consider the fraction *f*(*x*, *t*) of blue cells along a uniform, one-dimensional frontier at position *x* and time *t*. Every generation time *τ_g_*, the fraction *f*(*x*,*t*) will change due to the conversion probability *μ* and the competition at the frontier (which will depend on the selection coefficient s). For small *s* and *μ*, the fraction *f*(*x*,*t*) will evolve according to the stochastic differential equation of the stepping stone model (11):

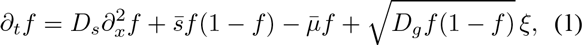

where 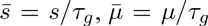 and ξ = ξ(*x,t*) is a Gaussian, white spatio-temporal noise with zero mean, 〈ξ(*x, t*)〉 = 0, and unit variance: 〈 ξ(*x,t*)ξ(*x’, t’*)〉 = *δ*(*t’ — t*)*δ*(*x’ — x*). The noise should be interpreted in the Ito sense (23), and describes the stochastic birth-death processes of the cells at the frontier, which have some effective genetic drift strength *Dg*. We expect the scaling *D_g_* ∽ *i/Nτ_g_*(11), where *l* is the linear size of the frontier over which cells compete to divide into virgin territory, and *N* is the number of these competing cells. We expect *t* to be a few cell diameters. The diffusion term 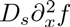 describes cell rearrangements caused by cell divisions at the frontier with an effective spatial diffusion constant *D_s_* ∽*l^2^*/*τ_g_*. The parameters *D_s_* and *D_g_* depend on the details of the microbial colony structure. They are measured for various microbial colonies in Refs. (12, 21). We will be primarily interested in how various solutions to Eq. 1 depend on *μ* and *s*, which we can control in the experiment.

Equation 1 belongs to the directed percolation universality class and exhibits a line of non-equilibrium phase transitions as a function of *μ* and *s* (7). The transition line may be found using Eq. 1 by examining sectors of unconverted cells as in Fig. 1(b) (i.e., by using Eq. 1 to evolve an initial *f*(*x,t =*0) with a localized “spike” of blue cells at the origin), but a uniform initial condition also exhibits a phase transition along the same phase boundary (9, 24). In particular, if we start with all blue cells at the initial frontier (*f*(*x,t* = 0) = 1), the average fraction of blue cells 〈*f*(*x,t*)〉*_x_* (averaged over the noise ξ in Eq. 1 and over all positions *x* along the frontier), will approach a non-zero constant 〈*f*(*x,t*)〉*_x_* → *f_∞_* > 0 as *t* → ∞ in the active phase and 〈*f*(*x,t*)〉*_x_*→ 0 in the inactive phase. A phase diagram similar to the one illustrated in Fig. 1(b) may then be constructed.

The directed percolation phase transition occurs along a line given approximately by *μ* ≈ *As^2^* for the range expansions (9), compared to *μ* ∽ *s* for well-mixed populations, where *A* is a constant of proportionality that will depend on *D_s_* and *D_g_*. We expect the noise term *D_g_* to be important near the conversional meltdown transition. In the strong noise limit, we derive an approximation for *A* by mapping the sector boundaries to random walks. Then, assuming the sectors do not collide, we find (9, 16, 25): 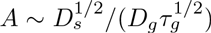. We may roughly estimate *A* by using measured values for the various parameters for a related yeast strain studied in Ref. (21): *D_s_* ≈ 15 *μ*m^2^/hr *, D_g_* ≈ 1.3 *μ*m/hr, and *τ_g_* ≈ 1.5 hr. With these estimates, we expect *A* ≈ 2. Note that the effective population size *N* ≈ 3 is quite small for these expansions, which is consistent with the evolutionary dynamics being dominated by competition at the very edge of the population. However, our growth conditions and yeast strains are different from Ref. (21), and a detailed check of the scaling of *A* with *D_s_* and *D_g_* is beyond the scope of this paper. Hence, we use *A* as a fitting parameter. Fitting *A* to our experimental results in Fig. 3 yields *A* ≈ 1.5, which is close to our crude estimate.

It is also possible to understand the sector angles illustrated in Fig. 1(a) using properties of the directed percolation universality class. First, note that a genetic sector formed from an unconverted (blue) cell at the frontier will have an opening angle *θ* given by (9)

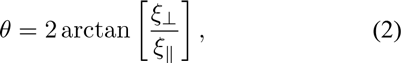

where ξ⊥/ξ_||_ is the slope of the sector boundaries, and ξ_||_ and ξ*⊥* are correlation lengths parallel and perpendicular to the growth direction. The opening angle can be measured in experiment. Note that long time sector survival requires we are in the active phase (see Fig. 1(b)), where there is a non-zero probability that the unconverted cell type will survive at long times. For each point (s, *μ*) in the phase diagram we define Δ = Δ(s, *μ*) as the shortest distance to the phase transition line. We will also change the sign of Δ as we cross the transition line, such that Δ > 0 in the active phase and Δ < 0 in the inactive phase. As we approach the phase transition line from the active phase (Δ → 0 with Δ > 0), we expect that the dynamics will be governed by the directed percolation phase transition (9). In particular, the slope ξ_⊥_/ξ _||_ of the sector (measured near the population frontier) is predicted to be proportional to a power of Δ:

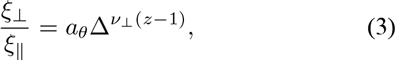

where *a_θ_* is a constant of proportionality, *z* ≈ 1.581 is a dynamical critical exponent, and *ν*_⊥__ ≈ 1.097 is a spatial correlation length exponent (24). The constant of proportionality *a_θ_* is not universal and will depend on the position along the transition line and on particular details of our model. So, as we approach the directed percolation transition, the sector angle *θ* is predicted to vanish according to

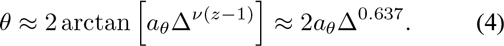

Unfortunately, the sector angle experimental results in Fig. 4 are too noisy to check the particular power-law behavior *θ ∽* Δ^0.637^ (although the data is consistent with this behavior). However, we can now check this particular power law prediction via range expansion simulations.

We simulate range expansions with flat, uniform frontiers (corresponding to a linear innoculation) on a triangular lattice with a single cell per lattice site. We take the frontiers of actively dividing cells to be a single cell wide and correspond to rows of the lattice. Cells at the frontier then compete with their neighbors to divide into the next lattice row. The probability of division is proportional to the cell growth rate. The unconverted blue cells have a growth rate normalized to 1, while the converted yellow cells grow with rate 1 — *s*. This protocol implements the selective advantage of the blue cells. After a cell division, the daughter cell mutates with probability *μ* if it is unconverted (just as in the designed yeast strain). These competition rules are a generalization of the Domany-Kinzel model updates (26). The lattice model is expected to be in the directed percolation universality class, as well. Our previous model, Eq. 1, is a possible coarse-grained description of the lattice model with *l* equal to the lattice spacing and an effective population size *N* = 1 (see Ref. (9) for details).

It is straightforward to evolve sectors by considering initial frontiers with just a single blue cell surrounded by all yellow cells. Some examples of the resulting sectors are shown in the insets of Fig. 1(b). The average angle *θ* subtended by the blue cell sectors is measured by calculating the width of a sector *W*(*t*), averaged over all sectors that survive to time *t*. Then, in the active phase, we expect *W*(*t*)≈ 2ξ_⊥_/ξ _||_t, from which we may extract the slope ξ⊥/ξ_||_ by fitting *W*(*t*) to a linear function. The angle is then extracted from Eq. 2 for various distances Δ away from the extinction transition line in the (*μ*, *s*) plane (see inset of Fig. 5). We find excellent agreement between simulation and Eq. 4 in Fig. 5. The parameter *a_θ_* ≈ 0.88 is found by fitting.

**Figure 5.**
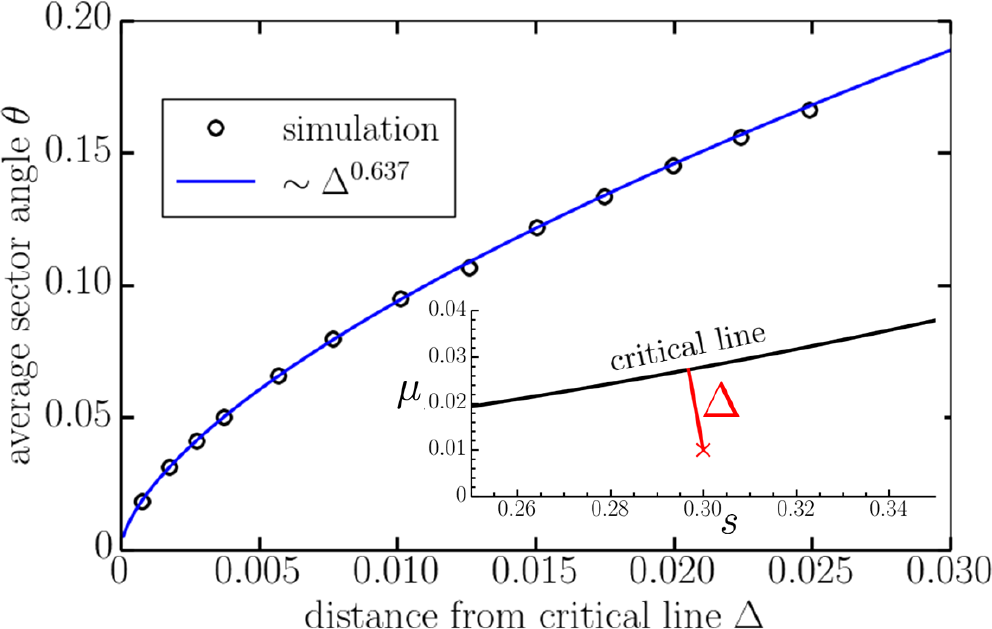
Average sector angles measured from 25600 simulations of two-dimensional range expansions as a function of the distance Δ away from the critical line separating the active and inactive phases shown in Fig. 1(b). The inset illustrates the shortest distance Δ to the critical line in the (s,*μ*)-plane. In the simulations, the distance Δ is varied by fixing *s =* 0.3 and varying the mutation rate *μ*. The range expansion has a flat frontier of 4000 cells and is evolved for 4 x 10^4^ generations. We initialize the populations with a single unconverted cell at the frontier and average the opening sector angle of all surviving sectors.

## Conclusion

We have examined an extinction transition using a genetically modified yeast strain that irreversibly converts from a more to a less fit strain. This synthetic strain maintains many sources of biological variability, including variability in growth rate and sector angle, while providing exquisite control over conversion, relative growth rate, and visualization of two cell types.

The experiments reveal that spatial dynamics enhances conversional meltdown, a major qualitative prediction of theory and simulations based on directed percolation ideas (9). Spatial fluctuations enhance extinction through genetic drift, which is much larger at population frontiers than in typical well-mixed experiments. According to theory, the extinction in a well-mixed population occurs when *μ* ∽ *s* and when *μ ∽ s^2^* for a range expansion with a thin (approximately one-dimensional) frontier. Hence, the unconverted strain is maintained in a smaller region of the (*μ*, *s*) phase space in the range expansion compared to the well-mixed case, as shown in Fig. 3. We expect that this enhancement is generic. The enhanced extinction probability might be observable in other spatially structured populations, such as tissue growth and natural range expansions. The enhanced extinction probability could have implications for maintaining stem cells populations and for cancer.

We also studied the opening sector angles of clusters of the fit strain spreading through a less fit population. In the flat front approximation, this opening angle is expected to vanish with a directed percolation power law as we approach the extinction transition (9). This power law was confirmed by simulations and is qualitatively consistent with experiments (8, 14). If front undulations are important, similar to systems described by the Kardar-Parisi-Zhang equation (27) or the noisy Burgers equation (28), the transition line discussed here in the context of directed percolation is expected to be in a different universality class. We then expect similar power law behavior, but with different critical exponents. Such power-law sector dynamics might be relevant for cancer, where driver mutations may spread through an otherwise slowly-growing cancerous population while accumulating irreversible, deleterious mutations (29). When many deleterious mutations can accumulate in parallel, we expect that there is an analogous extinction transition at which additional mutations accumulate fast enough to lead to a population collapse of the cells with the driver mutation (16, 29, 30).

To better understand these dynamics in models of precancerous tumors, we would need to consider three-dimensional range expansions with effectively two-dimensional frontiers, such as cells dividing at the surfaces of spherical masses of growing cells (31). In three-dimensional populations, the extinction dynamics could be quite different (30). Genetic drift is a weaker effect at two-dimensional frontiers, and the phase diagram for extinction will have a different shape. It would be interesting to examine three-dimensional range expansions of this synthetic strain to explore how these different spatial dynamics infu-ence the extinction transition. Experiments could be done by embedding the yeast in soft agar, or growing them up in cylindrical columns with nutrients supplied at the base, as described in Ref. (32).

## Acknowledgements

We thank Bryan Weinstein and Wolfram Moebius for helpful discussions. This work was supported by the National Science Foundation (NSF) through grant DMR-1306367, by the NIH through NIGMS grant GM068763, and by the Harvard Materials Research Science and Engineering Center (DMR-1420570). MOL also acknowledges support from NSF grant DMR-1262047. MEW was supported by the NDSEG and NSF GRFP fellowships. We thank Derek Lindstrom and Dan Gottschling for generously providing their inducible Cre construct P_*SCW11*_*cre-EBD78*. The computer simulations were run on the Odyssey cluster, maintained by the Harvard University Research Computing Group.

